# Autophagic activity acts as a rheostat in the control of nitrogen transfer from Arabidopsis rosette leaves to the seeds according to a principle of communicating vessels

**DOI:** 10.1101/2023.04.24.538060

**Authors:** Maxence James, Jacques Trouverie, Anne Marmagne, Fabien Chardon, Philippe Etienne, Céline Masclaux-Daubresse

## Abstract

Macroautophagy is known for long as essential for the degradation and the recycling of different macromolecules in eukaryotes. However how important is autophagy for nitrogen management at the whole plant level and for plant biomass and yield productivity in unstressed and well feed plants needed further investigation. In this study, we used both autophagy knock-out mutants and autophagy over-expressors that constitutively produce numerous autophagosomes. These mutants and over-expressors were cultivated using hydroponic system to observe and compare their phenotypes under sufficient nitrate supply, and when submitted after a while to strict nitrate starvation. The shift from nitrate sufficient condition to nitrate starvation allowed us to determine how autophagy defective or stimulated lines can use their own nitrogen resources to complete their cycle. Unexpectedly we observed that irrespective of the nitrate conditions, both mutants and over-expressors exhibited early leaf senescence phenotypes relative to wild type. While autophagy mutants exhibited strong defect for N remobilisation and seed production irrespective of nitrate condition, the better performance of autophagy-over expressors for N remobilisation and seeds production was only significant under sufficient nitrate supply, i.e. when autophagy was not naturally stimulated by nitrate limitation. Interestingly, comparisons of genotypes showed that the nitrogen pool used for seed filling originated from rosette leaves, as if rosette and seeds were used as communicating vessels independently of the stem and pod connecting organs. Altogether, results show that autophagy is a master player in nitrogen management at the whole plant level that controls yield production and leaf senescence.

## INTRODUCTION

Macroautophagy is a universal mechanism of eukaryotic cells that is essential for the degradation of unwanted cytoplasmic constituents. Macro-autophagy (here after named autophagy) consists in the formation of cytosolic double membrane vesicles named autophagosomes, that sequester cytoplasmic constituents dedicated to degradation, and drive them to lytic vacuoles in plant and yeast or to lysosomes in animal. Substrates for macroautophagy are diverse. They consist in damaged or useless proteins that have to be discarded, protein aggregates, portions of organelles and membranes. In the cell, autophagosomes act as “garbage trucks” that facilitates cell cleaning, ensure organelle quality control, and participates in the recycling of macromolecules in smaller molecules usable for cell nutrition or export. Autophagy is then essential for cell homeostasis and longevity in all eukaryotic organisms.

Autophagosome formation involves eighteen essential *AUTOPHAGY* (*ATG*) proteins that were first discovered in yeast (Tsukada and Ohsumi, 1993). Homologous genes were further identified in plant and animal (Yang and Bassham 2015). In Arabidopsis, several homologous yeast *ATG* genes belong to gene families. For example, there are nine *ATG8* genes in Arabidopsis (*ATG8a-i*). Several Arabidopsis *atg* mutants were studied since the discovery of *ATG* genes in plant. Mutants affected in single genes, like *atg5, atg7*, or *atg10* for example, display strong phenotypes as hypersensitivity to several abiotic stresses, reduced yield and plant biomass as well as early leaf yellowing (Phillips *et al*., 2008; Masclaux-Daubresse *et al*., 2014; Guiboileau *et al*., 2012; Guiboileau *et al*., 2013; Lenz *et al*., 2011; Yoshimoto *et al*., 2009; Thompson *et al*., 2005). The ATG8 proteins are the only proteins to be associated to the autophagosome membranes. The conjugation of ATG8 proteins with phosphatidylethanolamine (PE) is essential for autophagosome formation and the ATG8-PE conjugates that decorates the autophagosome membranes are involved in the recruitment and sequestration of the cytoplasmic material to be degraded (Noda *et al*., 2010; Masclaux-Daubresse *et al*., 2017). The *atg8* mutants do not display any specific phenotype relative to wild type. We then suppose that the different ATG8a-i proteins have redundant functions in targeting specific cargoes, and that redundancies explain the absence of phenotype for the *atg8* single mutants.

While abolishing autophagy through the mutation of *ATG* single genes is quite easy, stimulating autophagy genetically is more challenging as eighteen genes are involved in its core machinery. In yeast, a positive correlation between the amount of the ScATG8 in *Saccharomyces cerevisiae* cells and the size of autophagosomes and autophagy activity was found (Jin and Klionsky, 2014). It was postulated that the presence of more ATG8-PE conjugates in the cell could modulate membrane curvature and increase the size of the autophagosomes. In Arabidopsis, the overexpression of several *ATG8* genes under the control of constitutive promoters increased the number of autophagosomal structures (Chen *et al*., 2019). In rice, several studies confirmed that increasing the expression of *OsATG8* stimulate autophagy (Li *et al*., 2015).

In plant, like in other organisms, autophagy exists at a basal level in all the cells where it controls homeostasis. During senescence or in response to stresses, autophagy is enhanced to ensure organelle quality control and to discard oxidized material (Tang and Bassham, 2018). Nitrogen remobilization from senescing leaves is known to be an important process for plant yield and seed quality (Masclaux-Daubresse *et al*., 2010). The pioneer works of Guiboileau *et al*., (2012; 2013) showed that autophagy mutants display strong defects for protein degradation and nitrogen recycling in the vegetative tissues that resulted in a large decrease of the amount of organic nitrogen remobilized to the seeds during the seed filling period. Then, at the same time as autophagy is enhanced in senescing tissue to discard damaged cytoplasmic material, it contributes to recycling and export of nutrient from senescing tissues to new growing tissues. Chen *et al*., (2019) showed that constitutively over-expressing *ATG8* genes in Arabidopsis over-stimulates autophagic activity. Using ^15^N isotope, authors showed that N fluxes to the seeds were increased. Results from Guiboileau *et al*., (2012) and Chen *et al*., (2019) thus highlighted that autophagy could control nitrogen fluxes at the whole plant level.

In the present study we aimed at comparing in a same experiment the performances of autophagy knock-out mutants (*atg5* and *atg7*), *ATG8* over-expressors and wild type for seed production and nitrogen allocation to the seeds. We also aimed at characterizing better the nature of the source organs where defects or stimulation of autophagy play important roles for N remobilization to the seeds. Finally, this work contributed to better characterize the relationships between autophagy, N-remobilization and leaf senescence, using autophagy mutants and over-expressors as working models.

## MATERIAL and METHODS

### Plant material and growth conditions

The *Arabidopsis thaliana* autophagy T-DNA mutants *atg5-2* (SAIL_129B07) and *atg7-2* (GK-655B06) have been described previously Guiboileau *et al*., (2012) and by Lai *et al*., (2011). The *ATG8* over-expressors (*ATG8a*-OE and *ATG8g*-OE) have been constructed and described by Chen *et al*., (2019). Seeds were stratified for 48 h in 0.1 % agar (Select agar, Sigma, L’Isle d’Abeau Chesnes, France) at 4 °C in the dark and then sown onto lids cut from Eppendorf tubes (0.5 ml) and filled with 0.7 % agar. The plants of each genotype were placed in a glasshouse on a tank containing 10 L of 3.5mM NO_3_^-^ nutrient solution (1 mM KNO_3_, 1.25 mM Ca(NO_3_)_2_, 0.2 mM KH_2_PO_4_, 0.4 mM MgSO_4_, 0.1 mM K_2_SO_4,_ 50 μM NaFe-EDTA, 50 μM NaFe-EDDHA, 20 μM H_3_BO_3_, 6 μM MnSO_4_, 3 μM ZnSO_4_, 0.7 μM CuSO_4_, 0.008 μM (NH_4_)_6_Mo_7_O_24_, 0.1 μM CoCl_2_, 0.15 μM NiCl_2_, 0.9 mM Si(OH)_4_, 0.5 mM CaCl_2_, 0.1 mM KCl, 0.01 μM Na_2_Se0_4_, and 0.2 mM Na_2_SiO_3_ buffered to pH 6.8 with 0.36 mM CaCO_3_.) renewed weekly. Plants were grown on this solution for 44 days. At 44 days after sowing (DAS), plants were transferred to two contrasting N conditions: High Nitrate (HN; 3.5 mM nitrate) and No Nitrate (0N; 0.048 μM nitrate). The photosynthetic photon flux density was 110 mmol m^-2^ s^-1^ and day and night temperatures were 21 °C and 18 °C, respectively. Since sowing, plants were cultivated with a photoperiod of 8 h light/16 h dark. Then at 64 DAS, they were transferred on a 16h light/8h dark photoperiod to induce flowering and reproductive stage (Figure 1A).

**Figure 1:**
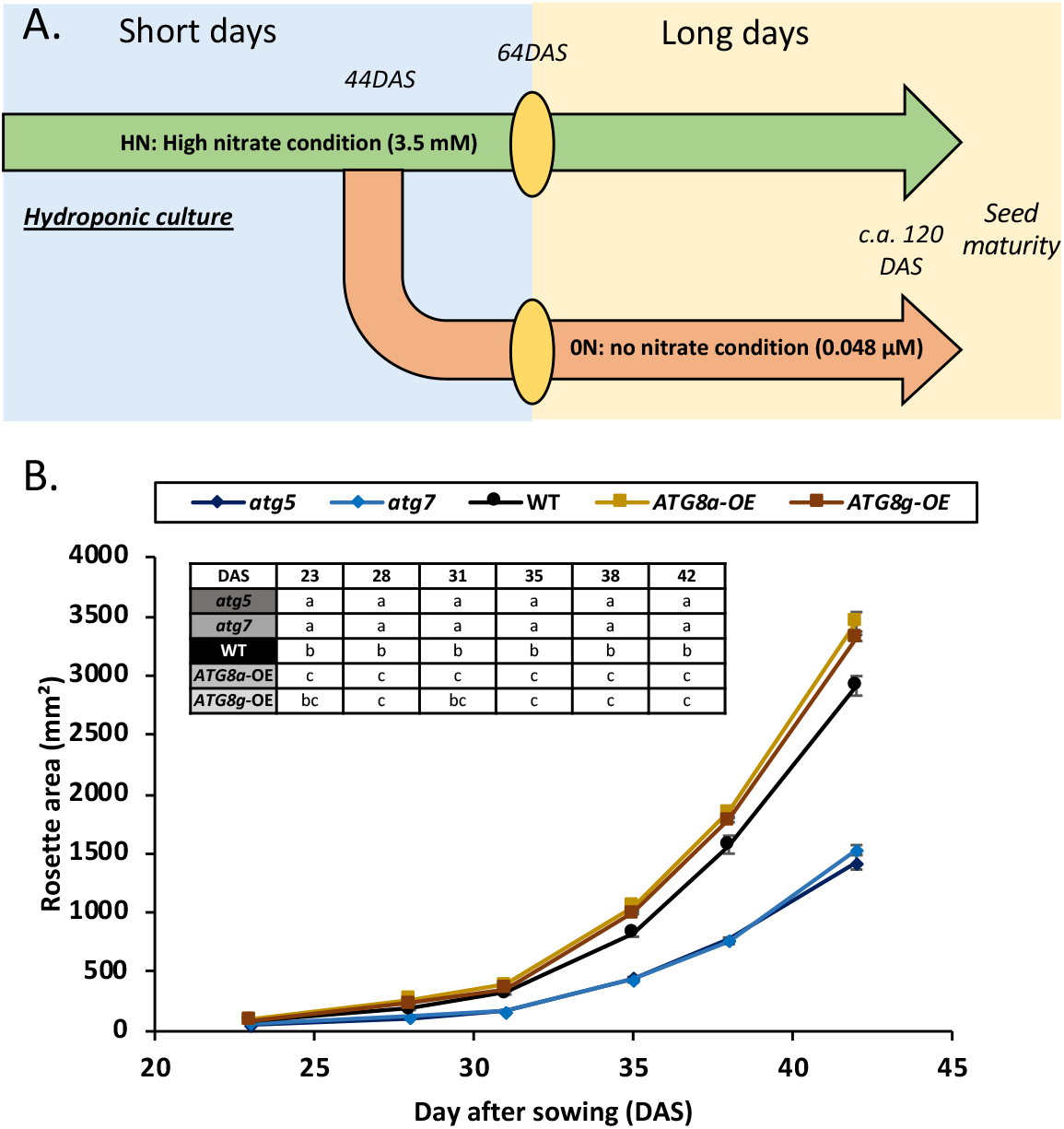
Leaf areas of WT, autophagy mutants (*atg5* and *atg7*) and over-expressors (*ATG8a*-OE and *ATG8g*-OE) reflect different growth under high nitrogen conditions. **(A)** Diagram of the experimental set-up for the study of autophagy mutants and over-expressors. The Col), autophagy mutants (*atg5* and *atg7*) and over-expressors (*ATG8a*-OE and *ATG8g*-OE) were cultivated on hydroponic system under nitrate (HN) condition in short days. Forty four days after sowing (DAS) half of the plants were transferred to no nitrate (0N). At 64 DAS all the plants were switched to long days. Harvesting time at seed maturity occurred at seed maturity. **(B)** Rosette areas were recorded once a week along the first 44 days of plant growth under HN. Values represent means ± SE for n≥11. Significant differences between types for each day after sowing are indicated by different letters in the table (p-value ≤ 0.05).

### Measurement of plant biomass and leaf area

Rosette areas were determined weekly during the first 44 DAS and leaf areas were measured on the separate leaf ranks at 64 DAS. The rosette and individual leaves were imaged (Canon Powershot G10, Canon, Japan, 14.8 Mpx, JPEG) to determine leaf areas using a specific script developed for ImageJ software (Schneider *et al*., 2012). The total area was determined with colour detection based on the HUE component of the Hue-Saturation-Lightness (HSL) system. The senescent leaf area was obtained from the difference between the total leaf area and the non-senescent leaf area corresponding of green area previously measured with the script.

At seed maturity, plants were harvested and stored in an oven (60 °C, 4 days) to obtain dry matter. The different organs were separated and their dry weights determined. Harvest index was calculated as the ratio between seed biomass and total plant biomass.

### Elemental analysis for C and N determination

At the seed maturity, N and C concentrations (as mg. 100 mg DW^-1^) were determined in the different compartment (Roots, leaves, stems, pods and seeds) of whole plant with an elemental analyser (EA3000, EuroVector, Milan, Italy). Nitrogen harvest index was then calculated as the ratio between the quantity of N in seeds and the total quantity of N in the whole plant.

### Statistical analysis

For all parameters, at least three biological replicates were measured (n ≥ 3). All data are presented as mean ± standard error (SE). To compare different data between different genotype or treatments, Tukey tests were performed after verifying compliance of normality with Shapiro-wilk test with R software. When the data did not conform to the normal distribution, they were transformed into Log. Statistical significance was postulated at p ≤ 0.05.

## RESULTS

### Both autophagy mutants and *ATG8* over-expressors exhibit more severe leaf senescence phenotypes than wild type

Rosette areas were recorded weekly for 44 days after sowing (DAS) on autophagy mutant (*atg5* and *atg7*), *ATG8* over-expressor (*ATG8a*-OE and *ATG8g*-OE) and wild type (WT) plants cultivated in hydroponic system with sufficient nitrate supply (HN; Figure 1A). From 28 DAS, rosette areas were significantly higher in *ATG8a-*OE and *ATG8g-*OE than in WT, and significantly lower in *atg* mutants compared to WT (Figure 1B) showing that defects in autophagy machinery impaired plant growth, while stimulation of autophagic activity improved plant growth and biomass under sufficient nitrate supply. This result was the first evidence of the positive role of enhanced autophagy activity on plant growth in unstressed environmental condition.

With plant ageing, the first symptoms of leaf senescence were observed on the rosettes of autophagy mutant (*atg5* and *atg7*) and *ATG8* over-expressor (*ATG8a*-OE and *ATG8g*-OE) after 60 DAS. Leaf senescence was then estimated measuring yellow and green areas on the fifteen first leaves of the rosettes at 64 DAS, i.e. on plants that have been transferred to sufficient (HN) or no (0N) nitrate conditions for 20 days (Figure 2AB). The proportions of yellow and green areas were compared between genotypes (Figure 2CD). Results show that irrespective of the nitrate condition, both mutants and over-expressors displayed larger yellowing symptoms than WT. Under HN, the proportion of yellowing area was three-times and two-times higher in autophagy mutants and *ATG8* over-expressors respectively, compared to WT. Under 0N, these differences between WT, mutants and over-expressors were lower than under HN, due to the fact that nitrate limitation is a strong senescence enhancing factor that attenuated genotype effects (Figures 2AB). These results show that both abolishment and stimulation of autophagy activity can results in early or more severe leaf senescence, which is in good accordance with the idea that fine tuning of autophagy is essential for longevity. Absence of macroautophagy may enhance cell death through intoxication from the accumulation of unwanted cell material, while excessive self-eating activity due to over stimulated autophagy could exhaust cell and drive them to cell (Guiboileau *et al*., 2010; Masclaux-Daubresse *et al*., 2017).

**Fig. 2.**
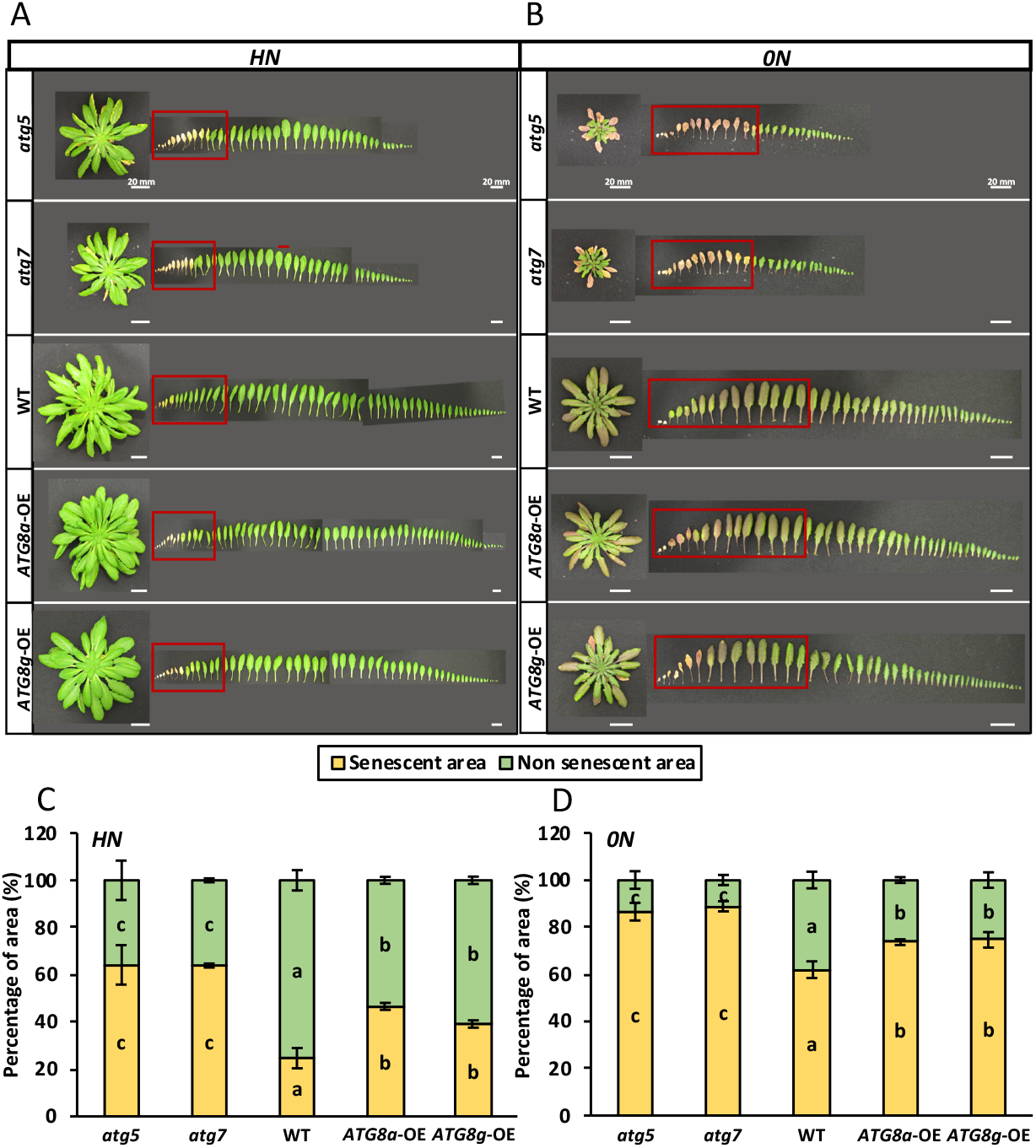
Autophagy mutants and over-expressors display higher leaf senescence symptoms than wild type. Leaves were dissected from WT, autophagy mutants (*atg5* and *atg7*) and overexpressors (*ATG8a*-OE and *ATG8g*-OE) transferred to high (HN; A, C) or no (0N; B, D) nitrate conditions for 20 days (64 DAS). Representative pictures of whole rosette, and leaf distribution from the base to the apex are presented (A, B). Histograms represent leaf areas split into senescent (yellow) and non-senescent (green) areas (C, D). The red squares in the pictures (A, B) correspond to the fifteen oldest leaves used to measure the percentages of senescent and non-senescent leaf areas and white bars correspond to 20 mm scale. The measures were performed on the fifteen oldest leaves dissected from four plant replicates for each genotype. All analyses were performed at 64 days after sowing (DAS). Values represent means ± SE for n=4. Significant differences between genotypes for each nitrogen condition are indicated by different letters (p-value ≤ 0.05).

### Autophagy controls seed yield production

Irrespective of the nitrate conditions, both *atg5* and *atg7* autophagy mutants displayed strong decrease in growth under hydropony. Their plant biomass at seed maturity was a fifth of that of WT (Fig. 3AB). Regarding the biomass partitioning, we can see that main developmental defect in autophagy mutant was the production of reproductive organs (stem, pods and seeds), (Fig. 3CD). Harvest index (HI), which is the ratio between seed yield and the total plant biomass, is an indicator of plant performance in terms of seed production per plant. Under both HN and 0N, autophagy mutants displayed significantly lower HI than WT (Fig. 4).

**Fig. 3.**
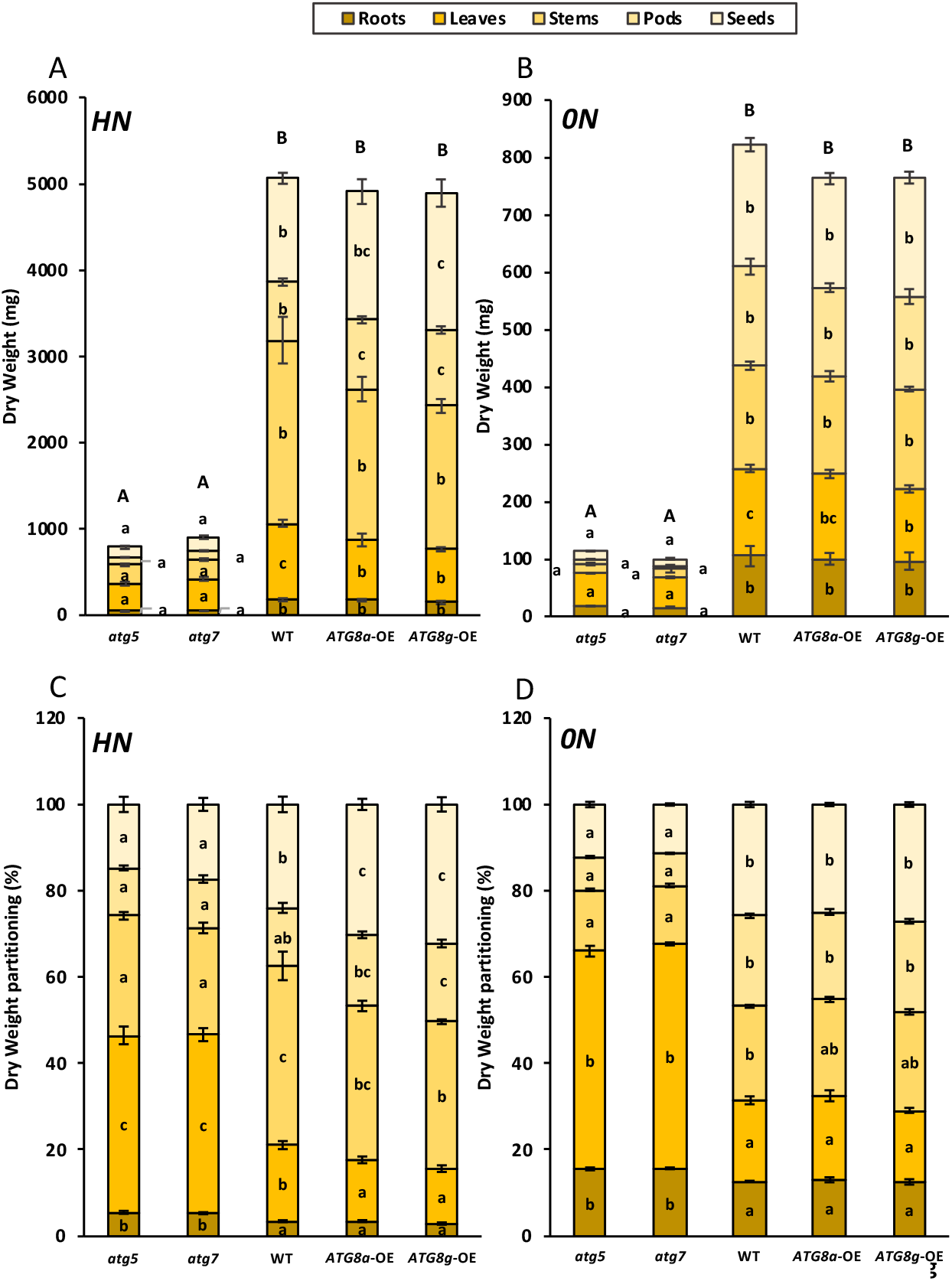
Dry weights and dry matter partitioning are different in both autophagy mutants and over-expressors compared to wild type. The dry weights of the different plant organs were recorded on plants harvested at seed maturity (A, B). The partitioning of dry matter in the different organs was calculated from dry weights and expressed as a % of the whole plant (C, D). WT, autophagy mutants (*atg5* and *atg7*) and overexpressors (*ATG8a*-OE and *ATG8g*-OE) plants were cultivated under high (HN; A, C) or no (0N; B, D) nitrate conditions. Values represent means ± SE for n=5. Significant differences between genotypes for each nitrogen condition are indicated by different letters (p-value ≤ 0.05).

**Fig. 4.**
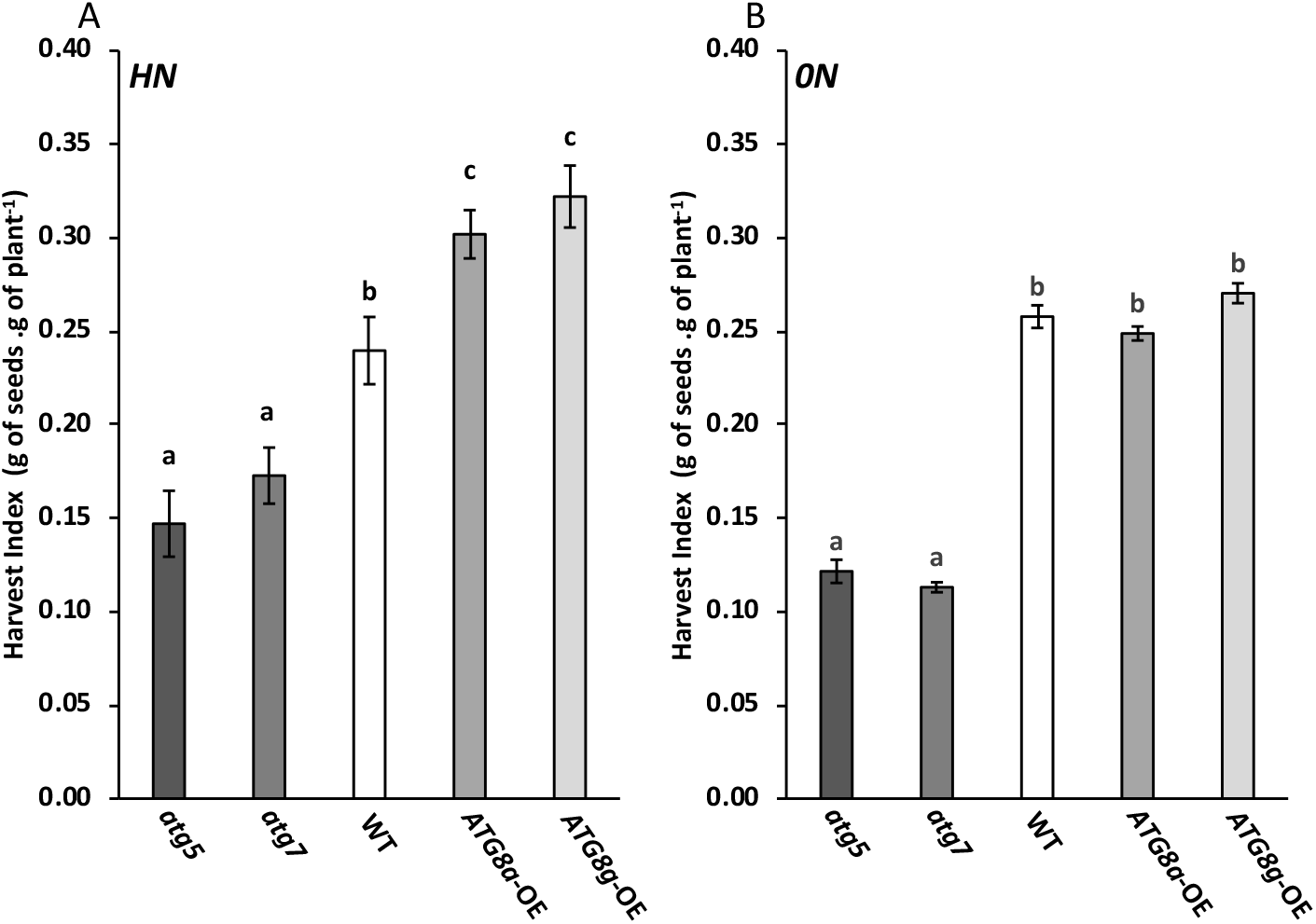
Harvest index are impacted in both autophagy mutants and over-expressors compared to wild type. The harvest index of the different plants were calculated on plants harvested at seed maturity (A, B). WT, autophagy mutants (*atg5* and *atg7*) and overexpressors (*ATG8a*-OE and *ATG8g*-OE) plants were cultivated under high (HN; A) or no (0N; B) nitrate conditions. Values represent means ± SE for n=5. Significant differences between genotypes for each nitrogen condition are indicated by different letters (p-value ≤ 0.05).

The total plant biomasses of the *ATG8a*-OE and *ATG8g*-OE over-expressors were not different from WT whatever HN or 0N (Fig. 3AB). However, under HN, leaf dry weights of *ATG8a*-OE and *ATG8g*-OE were lower than that of WT, while pod dry weights were higher (Fig. 3A). In addition, seed yield was significantly higher in *ATG8g*-OE and tends to be higher in *ATG8a*-OE than in WT under HN. Accordingly, biomass allocation in leaves and seeds was different in *ATG8a*-OE and *ATG8g*-OE compared to WT under HN (Fig. 3C), and *ATG8* over-expressors displayed higher HI than WT under HN (Fig. 4A). No difference between WT and the two *ATG8*-OE could be detected under 0N for both dry weights and biomass partitioning (Fig. 3 and 4).

Altogether these results reflect the better performance of the two *ATG8*-OE compared to WT when nitrate supply is sufficient. The absence of difference between WT and *ATG8*-OE under 0N can be explained by the fact that autophagic activity is primarily enhanced by nitrate limitation in all these genotypes and thus its over-stimulation by *ATG8* over-expression is inefficient. Thus, over-stimulating autophagy is a technical solution to increase yield under nitrate fertilizer supply.

### Nitrogen concentrations are strongly increased in all the organs of autophagy mutants

Autophagy *atg5* and *atg7* mutants, whose plant and organ biomasses are sharply lower compared to WT, displayed significantly higher nitrogen concentrations (N%) in all their organs (roots, leaves, stems, pods and seeds) under 0N (Fig. 5B), and also in roots, leaves and stems under HN (Fig. 5A). Although significant under 0N, the slight difference of N% between the *atg* mutants and WT pods and seeds reflects lower N allocation to the seeds in the *atg5* and *atg7* mutants which is consistent with the lower nitrogen remobilization efficiency of autophagy mutants previously reported by Guiboileau et al. (2012). The measured N concentrations suggest here that the defect of *atg* mutants for N remobilization mainly originates from roots, leaves and stems. The fact that N concentration was slightly higher in the seeds and pods of *atg* mutants, under 0N only, may be linked to the fact that seed production of autophagy mutants was especially affected under such conditions, leading then to less seeds but with higher N concentration.

**Fig. 5.**
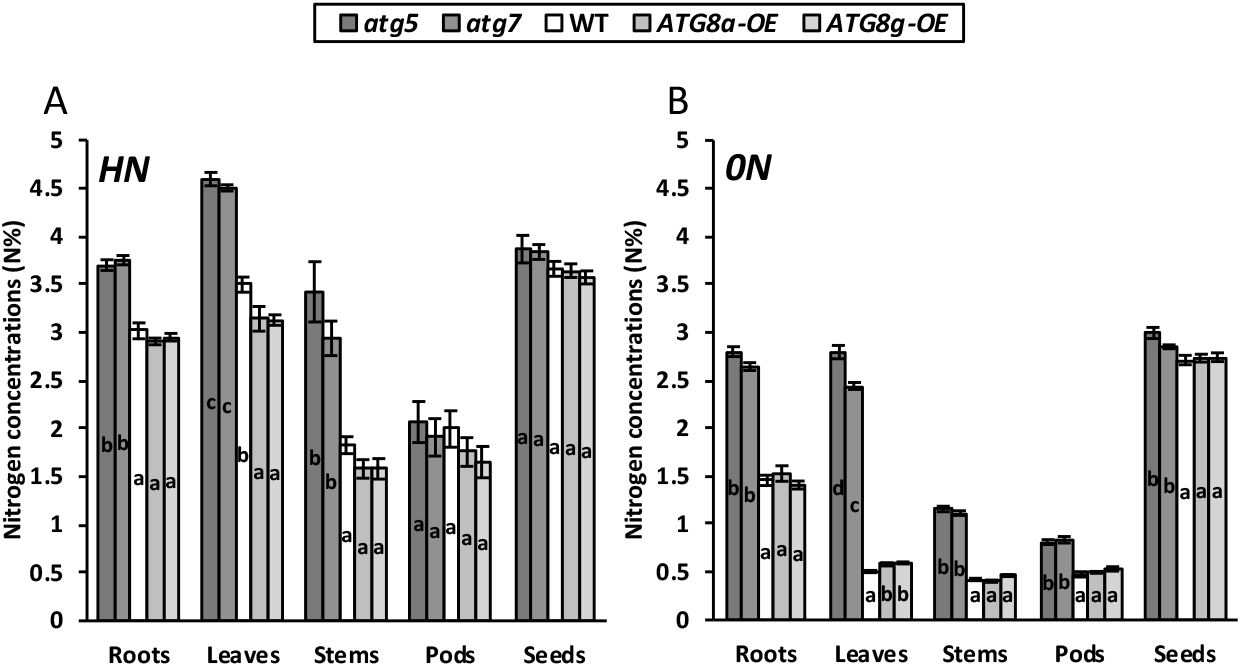
Higher nitrogen concentrations in the roots, stems and leaves of autophagy mutants reflect defects in nitrogen remobilisation to reproductive organs. The nitrogen concentrations in the different organs (roots, leaves, stems, pods and seeds) of WT, autophagy mutants (*atg5* and *atg7*) and over-expressors (*ATG8a*-OE and *ATG8g*-OE) were determined under high (HN; A) or no (0N; B) nitrate conditions. Nitrogen concentration were measured on plants harvested at seed maturity. Values represent means ± SE for n=5. Significant differences between genotypes for each nitrogen condition are indicated by different letters (p-value ≤ 0.05).

Differences in N concentrations between WT and *ATG8-OE* were minor and only detected in the leaves. N concentration was slightly lower in the leaves of *ATG8a*-OE and *ATG8g*-OE compared to WT under HN, but slightly higher under 0N (Fig. 5). The lower N% in the rosettes of *ATG8*-OE worth to be noticed as it indicates that less nitrogen remained in the vegetative parts of these plants at harvest, which is a mark of a better nitrogen use efficiency.

### Mutation and over-expression of autophagy gene trigger opposite effects on nitrogen allocation in leaves and seeds, reflecting inhibition and stimulation of source to sink N remobilization

N partitioning in the different organs was calculated from N concentrations and dry weights. Figure 6 shows that N partitioning in the seeds (i.e. nitrogen harvest index; NHI) and in the rosette leaves is strongly dependent of autophagy activity. Partitioning of N in the seeds of *atg* mutants is half and a third of that in seeds of WT under HN and 0N respectively (Fig. 7). The lower NHI of *atg* mutants is explained by a higher partitioning of N in their leaves, stems and roots, which are the main sources for N remobilization (Fig. 6). The discrepancies between *atg* mutants and WT in N partitioning are exacerbated under 0N condition, due to fact that the growth of stems, pods and seeds under 0N only relied on N remobilization as there was no more nitrate for root uptake.

**Fig. 6.**
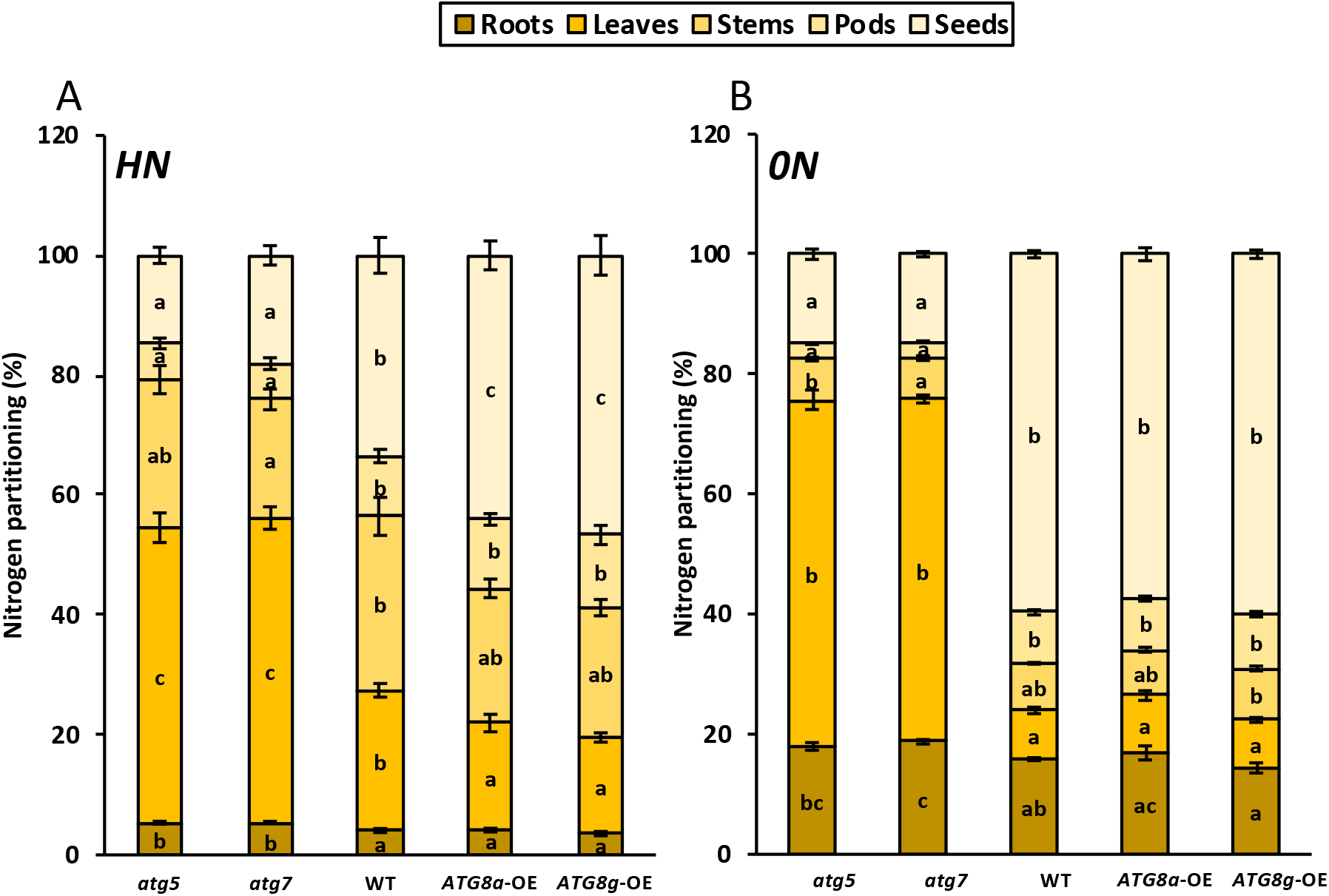
Autophagy activity is correlated to N remobilization from leaves to seeds in Arabidopsis under HN. Nitrogen partitioning (C, D) in the different organs of WT, autophagy mutants (*atg5* and *atg7*) and overexpressors (*ATG8a*-OE and *ATG8g*-OE) was calculated from nitrogen concentrations (Figure 4) and expressed as % of the whole plant. The nitrogen concentrations of the different compartments (roots, leaves, stems, pods and seeds) were measured on plants cultivated under high (HN; A) or no (0N; B) nitrate conditions, and harvested at seed maturity. Values represent means ± SE for n=5. Significant differences between genotypes for each nitrogen condition are indicated by different letters (p-value ≤ 0.05).

**Fig. 7.**
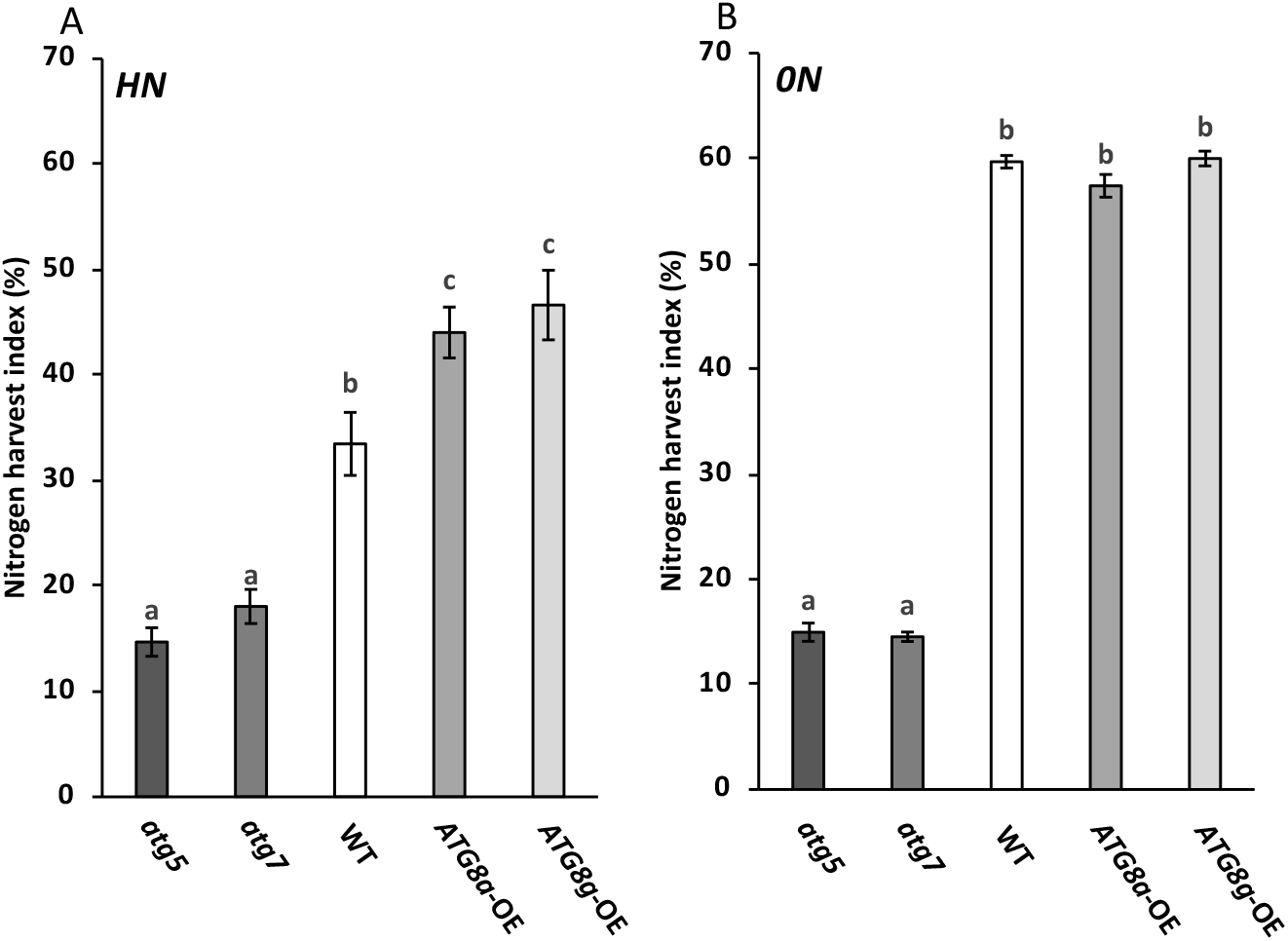
Nitrogen harvest index are lower in both autophagy mutants whatever the nitrate condition and higher in over-expressors specifically in high nitrate condition compared to wild type. The nitrogen harvest index of the different plants were calculated on plants harvested at seed maturity (A, B). WT, autophagy mutants (*atg5* and *atg7*) and overexpressors (*ATG8a*-OE and *ATG8g*-OE) plants were cultivated under high (HN; A) or no (0N; B) nitrate conditions. Values represent means ± SE for n=5. Significant differences between genotypes for each nitrogen condition are indicated by different letters (p-value ≤ 0.05).

No difference of N partitioning could be detected between the two *ATG8-OE* and WT under 0N. As said before, this was certainly due to the fact that autophagy activity was already enhanced to high level under 0N. By contrast, NHI was significantly higher in than in WT under HN (Fig. 7A), and such higher NHI was correlated to a lower N partitioning in the leaves of the *ATG8*-OE. Figure 6A nicely illustrates the opposite effects of autophagy stimulation (*ATG8*-OE) and autophagy inhibition (*atg*) on nitrogen allocation in source leaves and sink seeds.

Altogether these results show that the better NHI in autophagy over-expressors and lower NHI in autophagy mutant, result from the control of N remobilisation by autophagy in the rosettes leaves of Arabidopsis.

## DISCUSSION

The nitrogen budget study presented here confirms that autophagy activity is essential for N remobilization (as shown under 0N) and seed yield (Guiboileau *et al*., 2012). We also confirm that constitutively stimulating autophagy provides a strong benefice to plants, especially when nitrate supply is sufficient, increasing significantly seed yield and decreasing N concentrations in the dry rosette leaves at harvest. Increase in seed yield in the *ATG8-*OE under HN certainly relies on higher nitrogen remobilization as shown by Chen *et al*. (2019).

The dissection of plant organs permitted the determination of N partitioning in root, rosette, stems, pods and seeds at harvest. The assessment of the effect of modulating autophagy on the distribution of nitrogen in the plant suggests that autophagy acts as a tap controlling N partitioning between rosettes and seeds according to the principle of the communicating vessels (Figure 6; Figure 8). It was as if the stems and pods that make the physical link between rosette and seeds were not determinant in the control of nitrogen fluxes by autophagy.

**Fig. 8.**
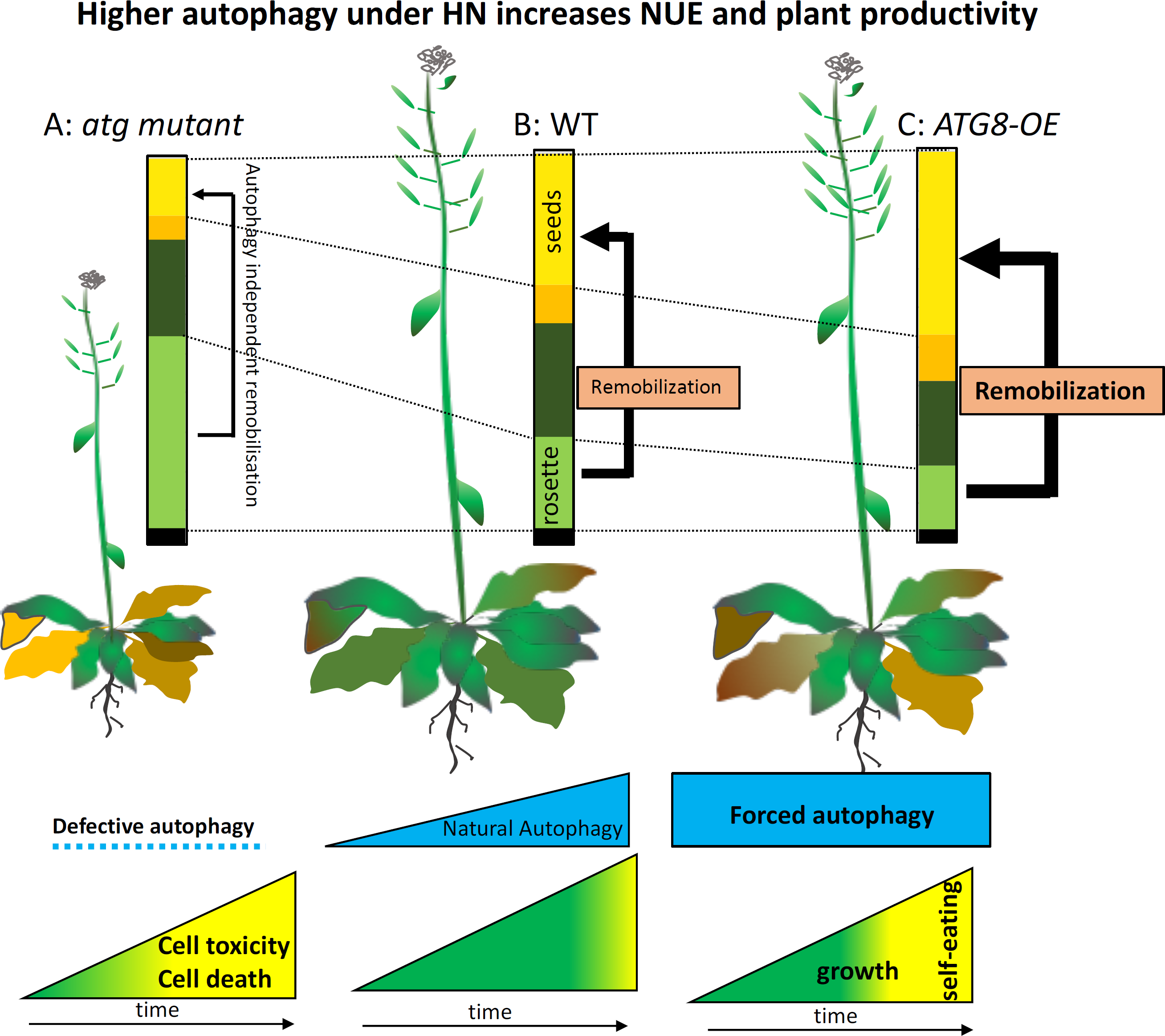
Schematic overview of the role of autophagy as a rheostat in nitrogen remobilization and leaf senescence under high nitrogen condition. The different situations, illustrated in blue, correspond to no autophagy (autophagy mutants; A), natural autophagy (WT; B) and ive autophagy (*ATG8* over-expressor; C). Histograms represent the nitrogen repartition in the different organs (from bottom to top: ack), rosette light green), stems (dark green), pods (orange), seeds (yellow)). Proportions are faithful to those presented in Fig. 6A. w and green in the different triangles at the bottom correspond to the proportion of senescent and non-senescent areas of the uring the vegetative stage according to the level of autophagy activity. Black arrows indicate nitrogen fluxes.

Other mechanisms than autophagy may then play a more prominent role in the regulation of nitrogen fluxes at the pod and stem levels. For example, Moison *et al*., (2018) showed that the cytosolic glutamine synthetases that are highly expressed in the phloem companion cells control N remobilization from the stem to the seeds. Such results were in good accordance with the fact that stems are the master organ for vascular tissues and sap circulation, and with the fact that glutamine synthetases are located in phloem tissues.

Literature dealing with N remobilization is largely related to leaf senescence (Havé *et al*., 2017*b*; Avice and Etienne, 2014; Gregersen *et al*., 2008; Brouquisse *et al*., 2000; Mae and Ohira, 1981). The temporal coincidence between leaf senescence and N remobilization metabolism has been so much reported that it is commonly accepted that nutrient remobilization is triggered by senescence, and that the role of leaf senescence is for nutrient remobilization. The phenotypes developed by autophagy mutants and over-expressors in our study underline the broader complexity of this association. Indeed, we show here that both autophagy mutants and over-expressors exhibit early yellowing phenotypes compared to WT, while they display opposite effects on N-remobilization efficiency relative to WT (Figure 8). This is certainly due to the fact that leaf longevity can control N remobilization and *vice-versa*. In *ATG8* over-expressors, the stimulation of the autophagic activity may improve cell cleaning thus cell longevity, at least for a while, permitting better recycling and nutrient export, which after a certain threshold certainly exhaust the cells and finally induces early senescence phenotype. At the reverse, in the absence of N remobilization due to autophagy defect, leaf tissues that are still nutrient rich, start to senesce as the absence of cellular cleaning by autophagy induces toxicities. Thus, both excess or lack of nutrient remobilization can induce early leaf senescence symptoms compared to WT fine-tuned autophagy level (Figure 8). This shows that the fine tuning of nitrogen management controls leaf longevity. As autophagy is not controlling 100% of the nitrogen remobilization flux, as shown by N partitioning in autophagy mutants under the 0N condition (Fig. 6B), we can also conclude that other mechanisms than autophagy are involved in N remobilization, but they obviously cannot compensate autophagy defect (James *et al*., 2018; Havé *et al*., 2017*a*).

In conclusion, the comparison of the performances of autophagy mutants and autophagy over-expressors presented here provides the demonstration of the master role of autophagy in the performance of plants for yield and nitrogen allocation to the seeds. The fact that stimulation of autophagy provides such improvement on seed yield and allocation of nitrogen to the seeds, under conditions of sufficient nitrate supply, is an important outcome. Indeed, today agriculture use nitrate fertilizer input, which is necessary for yield production. Therefore, our results show that forcing autophagy activity in the current agricultural context is a good technical solution to over-stimulate nitrogen fluxes to the seeds, decrease the amount of nitrogen lost in the vegetative tissues, and improve yield production (Figure 8). Our work also shows that enhancing autophagy constitutively in plant does not trigger deleterious effects as it has been previously suggested (Guiboileau *et al*., 2010). Although we observed a premature leaf senescence in the *ATG8*-OE, such phenotype was not correlated to lower biomass, yield or any kind of performance lost. In contrary it was accompanied by a better N allocation to the seeds and better yield. Thus *ATG8*-OE are the demonstration that leaf senescence can be a trait associated to better NUE performance and better harvest index, as far as it does not kill too early plant tissue, like it is the case in autophagy mutants. The reason why this senescence trait is associated to positive performances in *ATG8-*OE is certainly due to the fact that autophagy, while involved in degrading process, is selective and thus guaranties healthy and well-functioning cells until they are nutrient exhausted and die. Therefore, finding technical solutions, and especially non-transgenic solutions, for autophagy induction is a valuable innovation to promote sustainable agriculture and healthy planet.

## ACKNOWLEDGMENTS

We are most grateful to PLATIN’ (Plateau d’Isotopie de Normandie) core facility for all element and isotope analysis used in this study and the technical staff of UMR 950 EVA (Magalie Bodereau, Josiane Pichon, Elise Nexer and Kenza Boulouiz) for their help in harvesting and dissecting the plants. The authors declare that the research was conducted in the absence of any commercial or financial relationships that could be construed as a potential conflict of interest.

## AUTHOR CONTRIBUTIONS

Designed research: PE, JT, AM, FC and CMD; conducted experiments: MJ; supervision of experiments: PE, JT; analyzed data: MJ and CMD; discussed results: MJ, PE, JT, AM, FC and CMD; wrote paper: MJ, CMD, PE

## CONFLICT OF INTEREST

No conflict of interest declared

## FUNDING

This work was funded by the French National Research Agency (ANR-19-CE14-0009-02 hAPPEN: Autophagy, Proteases and plant Performances). The IJPB benefits from the support of Saclay Plant Sciences-SPS (ANR-17-EUR-0007).

## Notes

### Competing Interest Statement

The authors have declared no competing interest.

## REFERENCES

Avice JC, Etienne P. 2014. Leaf senescence and nitrogen remobilization efficiency in oilseed rape (Brassica napus L.). Journal of Experimental Botany 65, 3813–3824.

Brouquisse R, Masclaux C, Feller U, Raymond P. 2000. Protein hydrolysis and nitrogen remobilisation in plant life and senescence. The assimilation of nitrogen by plants, 400p, in press.

Chen Q, Shinozaki D, Luo J, et al. 2019. Autophagy and nutrients management in plants. Cells, 1426.

Gregersen PL, Holm PB, Krupinska K. 2008. Leaf senescence and nutrient remobilisation in barley and wheat. Plant Biology 10, 37–49.

Guiboileau A, Avila-Ospina L, Yoshimoto K, Soulay F, Azzopardi M, Marmagne A, Lothier J, Masclaux-Daubresse C. 2013. Physiological and metabolic consequences of autophagy defisciency for the management of nitrogen and protein resources in Arabidopsis leaves depending on nitrate availability. New Phytologist 199, 683– 694.

Guiboileau A, Sormani R, Meyer C, Masclaux-Daubresse C. 2010. Senescence and death of plant organs: Nutrient recycling and developmental regulation. Comptes Rendus Biologies 333, 382–391.

Guiboileau A, Yoshimoto K, Soulay F, Bataillé MP, Avice JC, Masclaux-Daubresse C. 2012. Autophagy machinery controls nitrogen remobilization at the whole-plant level under both limiting and ample nitrate conditions in Arabidopsis. New Phytologist 194, 732–740.

Havé M, Balliau T, Cottyn-Boitte B, et al. 2017a. Increase of proteasome and papain-like cysteine protease activities in autophagy mutants: backup compensatory effect or pro cell-death effect? Journal of Experimental Botany 69, 1369–1385.

Havé M, Marmagne A, Chardon F, Masclaux-Daubresse C. 2017b. Nitrogen remobilization during leaf senescence: lessons from Arabidopsis to crops. Journal of Experimental Botany 68, 2513–2529.

James M, Poret M, Masclaux-Daubresse C, Marmagne A, Coquet L, Jouenne T, Chan P, Trouverie J, Etienne P. 2018. SAG12, a Major Cysteine Protease Involved in Nitrogen Allocation during Senescence for Seed Production in Arabidopsis thaliana. Plant and Cell Physiology 59, 2052–2063.

Jin MY, Klionsky DJ. 2014. Regulation of autophagy: Modulation of the size and number of autophagosomes. Febs Letters 588, 2457–2463.

Lai Z, Wang F, Zheng Z, Fan B, Chen Z. 2011. A critical role of autophagy in plant resistance to necrotrophic fungal pathogens. Plant Journal 66, 953–968.

Lenz HD, Haller E, Melzer E, et al. 2011. Autophagy differentially controls plant basal immunity to biotrophic and necrotrophic pathogens. Plant Journal 66, 818–830.

Li WW, Chen M, Zhong L, Liu JM, Xu ZS, Li LC, Zhou YB, Guo CH, Ma YZ. 2015. Overexpression of the autophagy-related gene SiATG8a from foxtail millet (Setaria italica L.) confers tolerance to both nitrogen starvation and drought stress in Arabidopsis. Biochemical and Biophysical Research Communications 468, 800– 806.

Mae T, Ohira K. 1981. The remobilisation of nitrogen related to leaf growth and senescence in rice plants (Oriza sativa L.). Plant and Cell Physiology 22, 1067–1074.

Masclaux-Daubresse C, Chen Q, Havé M. 2017. Regulation of nutrient recycling via autophagy. Current Opinion in Plant Biology 39, 8–17.

Masclaux-Daubresse C, Clément G, Anne P, Routaboul JM, Guiboileau A, Soulay F, Shirasu K, Yoshimoto K. 2014. Stitching together the multiple dimensions of autophagy using metabolomic and transcriptomic analyses reveals new impacts of autophagy defects on metabolism, development and plant response to environment. The Plant Cell 26, 1857–1877.

Masclaux-Daubresse C, Daniel-Vedele F, Dechorgnat J, Chardon F, Gaufichon L, Suzuki A. 2010. Nitrogen uptake, assimilation and remobilization in plants: challenges for sustainable and productive agriculture. Annals of Botany 105, 1141–1157.

Moison M, Marmagne A, Dinant S, et al. 2018. Three cytosolic glutamine synthetase isoforms localized in different-order veins act together for N remobilization and seed filling in Arabidopsis. Journal of Experimental Botany 69, 4379–4393.

Noda NN, Ohsumi Y, Inagaki F. 2010. Atg8-family interacting motif crucial for selective autophagy. Febs Letters 584, 1379–1385.

Phillips AR, Suttangkakul A, Vierstra RD. 2008. The ATG12-conjugating enzyme ATG10 is essential for autophagic vesicle formation in Arabidopsis thaliana. Genetics 178, 1339–1353.

Schneider CA, Rasband WS, Eliceiri KW. 2012. NIH Image to ImageJ: 25 years of image analysis. Nature Methods. 671–675.

Tang J, Bassham DC. 2018. Autophagy in crop plants: what’s new beyond Arabidopsis? Open Biology 8.

Thompson AR, Doelling JH, Suttangkakul A, Vierstra RD. 2005. Autophagic nutrient recycling in Arabidopsis directed by the ATG8 and ATG12 conjugation pathways. Plant Physiol 138, 2097–110.

Tsukada M, Ohsumi Y. 1993. Isolation and characterization of autophagy defective-mutants of Saccharomyces cerevisiae. Febs Letters 333, 169–174.

Yoshimoto K, Jikumaru Y, Kamiya Y, Kusano M, Consonni C, Panstruga R, Ohsumi Y, Shirasu K. 2009. Autophagy Negatively Regulates Cell Death by Controlling NPR1-Dependent Salicylic Acid Signaling during Senescence and the Innate Immune Response in Arabidopsis. Plant Cell 21, 2914–2927.

